# Predation on scyphozoan polyps and selective nematocyst incorporation dynamics in the aeolid, *Caloria militaris*

**DOI:** 10.1101/2024.12.29.630659

**Authors:** Hila Dror, Tamar Lotan, Dror Angel

## Abstract

Aeolid nudibranchs are known to prey on various cnidarians and incorporate nematocysts from their prey into their cerata (dorsal appendages) for self-defense. Nematocyst incorporation is selective, and is based on species, diet, prey availability, and predation pressure. This study investigated the interactions between two nudibranch species, *Caloria militaris* and *Flabellina affinis*, and scyphozoan polyps from common eastern Mediterranean jellyfish species, including *Rhopilema nomadica*, *Cassiopea andromeda*, *Phyllorhiza punctata*, and *Aurelia* sp. Laboratory experiments were used to assess whether these nudibranchs feed on scyphozoan polyps, and if they incorporate the prey nematocysts into the cerata. Results indicate that while *F. affinis* avoids scyphozoan polyps, *C. militaris* consumes all tested species, and can subsist on them for extended durations (up to 255 days). Predation rates varied among prey species tested, ranging from a mean of 8 to 43.3 polyps day^-1^. However, *C. militaris* does not store scyphozoan nematocysts. These findings contribute to understanding nudibranch feeding ecology and the potential roles these predators play in regulating jellyfish blooms. Identification of scyphozoan nematocysts in the cerata of predator aeolid nudibranchs is discussed as a means to locate cryptic scyphozoan polyps in the marine environment.

## 1. Introduction

Aeolid nudibranchs (Gastropoda: Opisthobranchia) are marine gastropods that prey on cnidarians such as jellyfish, anemones, and hydroids^1–5^. Many are highly specialized feeders, targeting specific prey, and their larvae settle where adult prey is abundant^2,3,6,7^. Nudibranchs have evolved defenses against the cnidarian stinging mechanism, which consists of syringe-like capsules known as nematocysts^8,9^. These defenses include protective gut linings, epithelia, and mucus secretion^1,10–14^. Through kleptocnidae, nudibranchs retain and use their prey’s nematocysts for defense, storing them in cnidosacs at the tips of their cerata (dorsal appendages)^10,12,15^. Not all nematocysts are retained, indicating selective sequestration^16–18^, which may vary with prey availability and predator presence, with some species preferring scyphozoan nematocysts when multiple prey types are available^19^.

The retained nematocysts may persist within the nudibranch’s body for extended periods, ranging from days to weeks. Since the inventory of nematocysts (termed cnidome) facilitates taxonomic identification of Cnidaria^20^, the presence and composition of nematocysts within the cerata can provide insights into the nudibranch’s dietary history, serving as a record of the past and more recently consumed prey^10,17,21,22^.

Here, we concentrated on two aeolid species, *Flabellina affinis* (Family: Flabellinidae) (Gmelin, 1791) (Fig. 1a), a native aeolid to the Mediterranean, and *Caloria militaris* (Family: Facelinidae) (Alder and Hancock, 1864) (Fig. 1b), a Lessepsian migrant^23^. *F. affinis*, measuring up to 40 mm in length, is present throughout the Mediterranean and along the Atlantic African coasts (http://www.seaslugforum.net/find/flabaffi). It is known to inhabit various habitats including seaweed beds, rocky reefs, and soft bottom, up to 10 meters deep, and was found to feed on hydroids of the genus *Eudendrium*^24^*. C. militaris* was originally described in the Bay of Bengal, India, and has been recorded throughout the Indo-West Pacific (http://www.seaslugforum.net/factsheet/phidmili). This species is known to inhabit natural hard substrates, although it can also be found on artificial substrates up to 30 meters deep. *C. militaris*, measuring up to 40 mm in length, is known to feed on hydroids such as *Calyptospadix cerulea* (Clarke, 1882), *Ectopleura larynx* (Ellis & Solander, 1786), and *Pennaria disticha* (Goldfuss, 1820)^3,25^. Along the Israeli Mediterranean coast, *C. militaris* is now commonly found in large numbers during winter and spring, often in association with the hydroid *P. disticha* (Dror, personal observation).

**Fig. 1:**
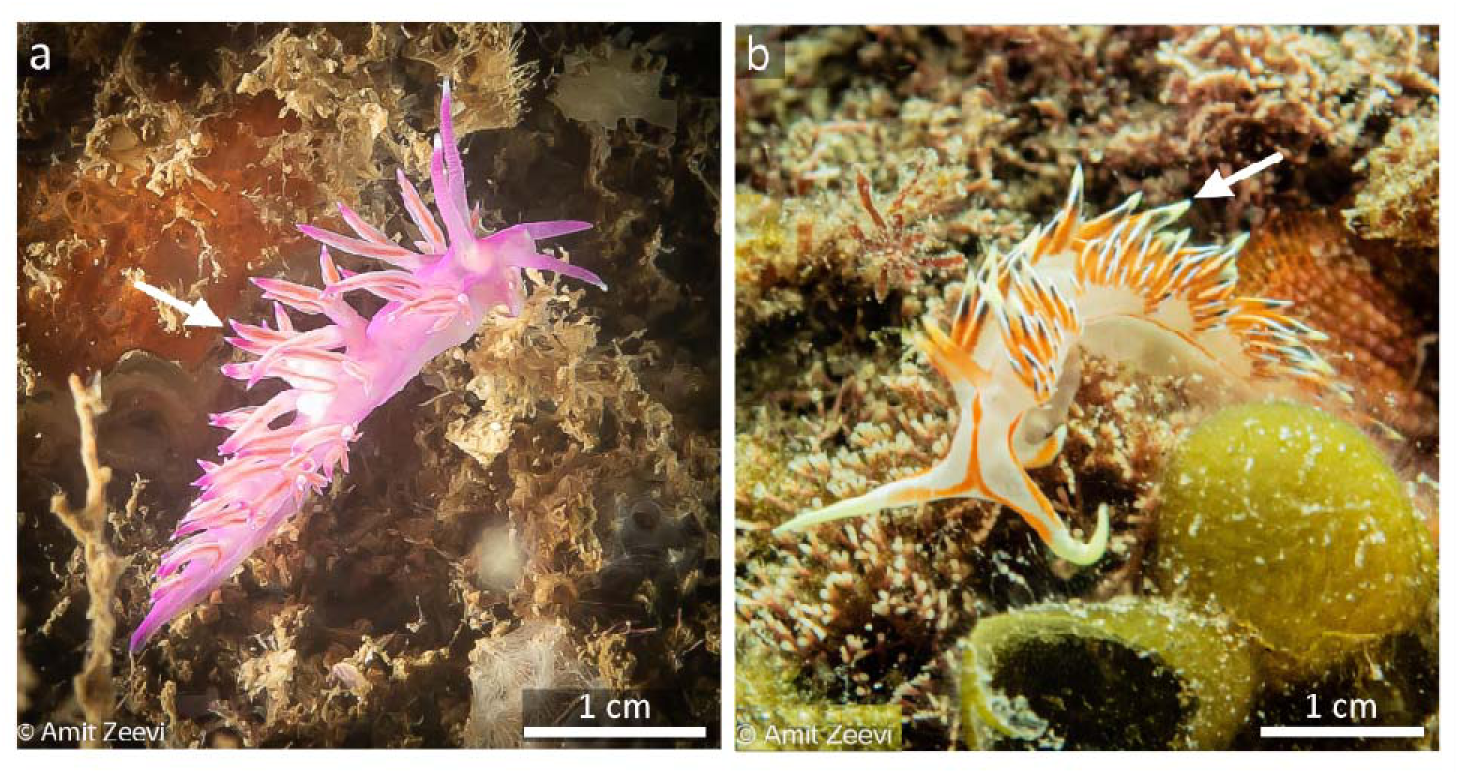
The aeolid nudibranchs (a) *F. affinis* and (b) *C. militaris* . Arrows mark the tips of the cerata. Photos: Amit Zeevi.

The eastern Mediterranean has been experiencing massive jellyfish blooms during the last several decades, which have negative ecological, economic and societal effects^26^. We wanted to explore the interactions between nudibranchs and Scyphozoa species common to the region. *Rhopilema nomadica* (Galil, Spanier & Ferguson, 1990), *Cassiopea andromeda* (Forskål, 1775), and *Phyllorhiza punctata* (von Lendenfeld, 1884) are Indo-Pacific species that occur in the eastern Mediterranean^27^, whereas *Aurelia* sp. is a cosmopolitan genus^28^. Since the 1980’s, *R. nomadica* has become the dominant jellyfish species in the eastern Mediterranean, with population present most of the year^29^. It is considered one of the “100 worst invasive species” in the Alien Invasive Species Inventories for Europe^30^ due to its negative ecological and economic effects and rapid expansion westward^26,31^.

Most Scyphozoa have a complex life cycle, consisting of both a pelagic medusa stage and a benthic polyp stage. Polyps are regarded as the driving force behind jellyfish outbreaks since they reproduce asexually and can persist for extended periods, releasing numerous young medusae each season^2,32^. The polyps of most of the Scyphozoa are cryptic and their location at sea remains unknown. Nudibranchs have been found to effectively consume scyphozoan polyps^2,33–36^, leading to a significant reduction in polyp populations^2^. Therefore, understanding the feeding behavior of nudibranchs and their predation on scyphozoan polyps is important in our quest to understand the dynamics of scyphozoan populations^2,33–36^.

We examined the interactions between aeolid nudibranchs and scyphozoan polyps, focusing on species commonly found in the eastern Mediterranean. First, we tested which nudibranch species can feed on scyphozoan polyps and whether these can serve as a primary food source for extended periods. We hypothesized that certain nudibranch species exhibit a more generalist feeding behavior than previously reported, capable of surviving solely on scyphozoan polyps when their preferred food (Hydrozoa) is scarce. Next, we examined whether these nudibranchs can incorporate the nematocysts of prey scyphozoan polyps into their cerata. We found that predation by *C. militaris* may serve as an important factor regulating scyphozoan polyp populations.

## 2. Results

### 2.1. Short-term predation of nudibranchs on various cnidarian

All nudibranchs that were placed in the bowls explored the bottom, sides, water surface, and the slide with polyps. However, *F. affinis* avoided contact with *R. nomadica* polyps, and no polyps were consumed during the 24 h experiment (Supp. Fig. S1). In contrast, *C. militaris* retracted when its oral tentacles first contacted the polyp, as if stung, and either moved away or remained still. To avoid being stung when feeding, the nudibranch elevated its head, and engulfed the polyp from the top, consuming the goblet and leaving the stalk (Supp. material S2). During the close-up observation, three out of the four *C. militaris* individuals were observed feeding on *R. nomadica* polyps. Twenty-four h later, between 58% and 100% of the polyps that were offered had been consumed by *C. militaris* nudibranchs (19 ± 8.83 polyps day^-1^). However, when *C. militaris* were offered adult *R. nomadica* tentacles, they kept their distance. Upon contact, the nudibranch would get stung, curl up, roll and quickly crawl away. Predation of *C. militaris* on *C. andromeda* polyps was less efficient than on *R. nomadica* polyps. Upon first contact, the polyp attacked the nudibranch causing it to retreat. The first *C. andromeda* polyp was approached only 82 min after *C. militaris* was added to the experimental bowl, and it took nearly 30 min for the nudibranch to fully swallow it. By the end of the 24 h experiment, the nudibranch had consumed only three *C. andromeda* polyps (37.5% predation). In contrast to the behavior of *C. militaris* offered scyphozoan polyps, when presented with the hydrozoan, *P. disticha*, it began feeding immediately, and continued throughout the observation period. The nudibranch would crawl on the hydrozoan branch, reach a polyp, remove it from the stalk, and swallow it whole. The sting of the polyps did not seem to affect the nudibranch as it didn’t retract when coming into contact with the polyp tentacles. Unlike the hydrozoan and scyphozoan polyps, none of the anemone (*E. diaphana*) and coral (*O. patagonica*) polyps were consumed by *C. militaris.* The nudibranch made some attempts to approach the anemone but was severely stung and kept away. Our findings indicate that scyphozoan and hydrozoan polyps may serve as prey for *C. militaris* nudibranchs in the wild.

### 2.2. Long-term predation of *C. militaris* on scyphozoan polyps

*C. militaris* nudibranchs were offered four species of scyphozoan polyps as prey. The mean (and maximum) number of polyps consumed by the nudibranchs per day was 8 ± 10.2, 32.8 ± 23.7, 44 ± 45.3, 43.3 ± 24.1 (49, 130, 192, and 105) for *Aurelia* sp., *C. andromeda*, *R. nomadica*, and *P. punctata,* respectively (Table 1). The predation rate and the predation proportion was significantly lower for the *Aurelia* sp. polyps compared to the other species (*C. andromeda*, *R. nomadica*, and *P. punctata* ) (F_(3,217)_ = 19.3, p < 0.0001) (Fig. 2). Whereas thousands of *R. nomadica* polyps were consumed during the experiment, the podocysts of *R. nomadica* were left untouched. Nudibranch survival was longest when feeding on *C. andromeda* polyps (146 - 255 days) and the shortest when feeding on *P. punctata* polyps (34 - 40 days) (Supp. video S2-S4: and Supp. Table S5). The nudibranchs started feeding almost immediately upon encounter with prey and no ingestive conditioning to a certain prey was observed (no change in predation rate over time). To better understand the potential effect of predation of *C. militaris* on scyphozoan polyps, we scanned the literature for nudibranch species known to feed on scyphozoan polyps (Table 1), and found that *C. militaris* predation rate is within the range of predation rates reported for other nudibranch species. These results indicate that *C. militaris* may be a prominent predator of scyphozoan polyps.

**Fig. 2:**
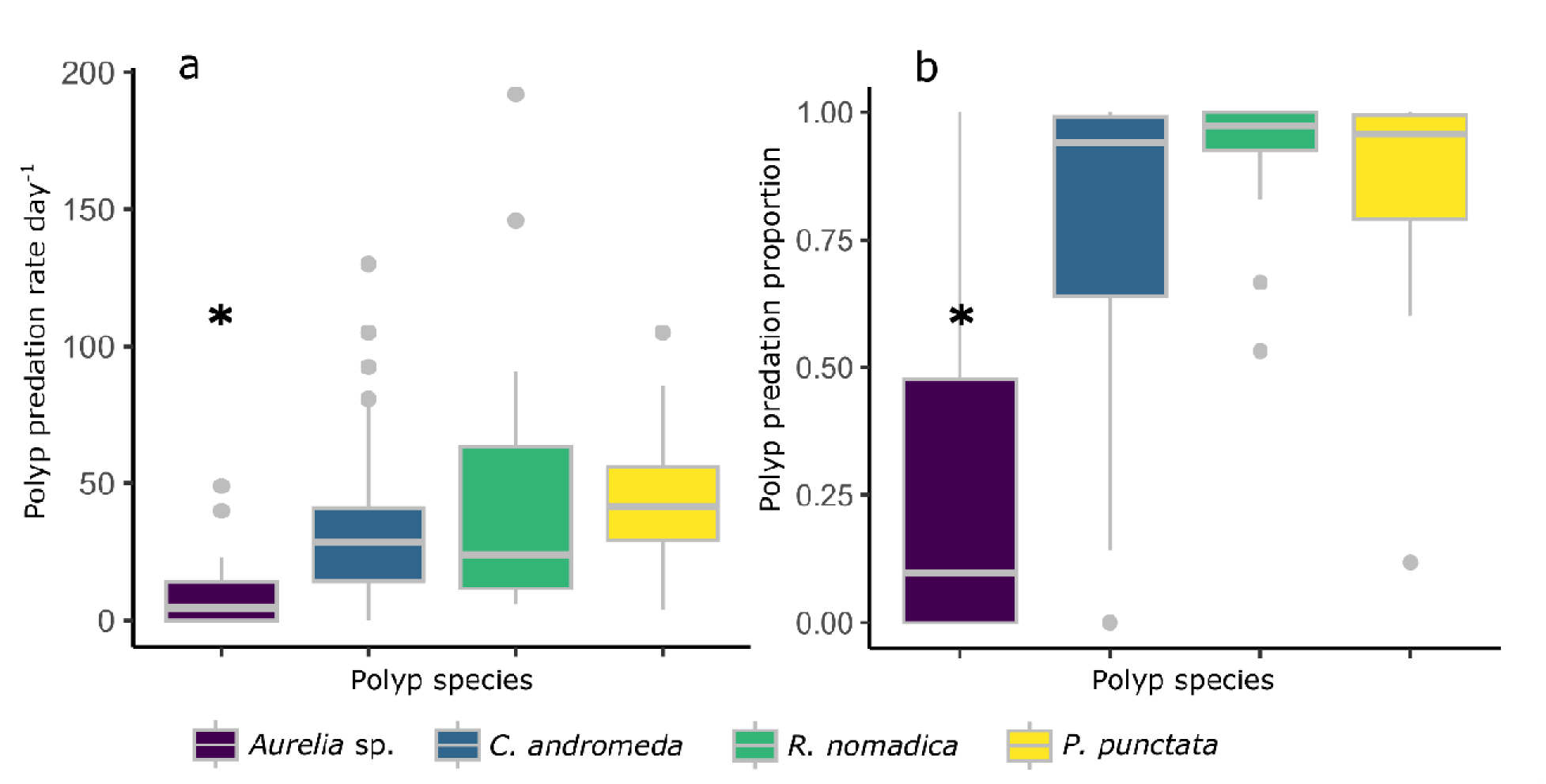
Predation of *C. militaris* on *Aurelia* sp., *C. andromeda* , *R. nomadica,* and *P. punctata* polyps. (a) Predation rate presented as polyps day^-1^ (b) Predation proportion. Asterisk (*) indicates significance at P < 0.0001. Number of nudibranchs tested for each prey species: 5, 4, 2, and 4 for *Aurelia* sp., *C. andromeda*, *R. nomadica,* and *P. punctata,* respectively.

**Table 1:** Nudibranch species that prey on scyphozoan polyps and predation rate (when available). * Predation rates are provided as polyps day^-1^, unless otherwise stated. ** Dependent on body size.

| Nudibranch | Schyphopolyp | Citation | Predation rate (polyps d<sup>-1</sup>) * |
| --- | --- | --- | --- |
| ***Caloria militaris***<br>(Alder & Hancock, 1864) | *Aurelia* sp. | This study | 8 ± 10.2 |
|  | *Cassiopea andromeda* | This study | 32.8 ± 23.7 |
|  | *Phyllorhiza punctata* | This study | 43.3 ± 24.1 |
|  | *Rhopilema nomadica* | This study | 44 ± 45.3 |
| ***Ceratodoris plana***<br>(as *Okenia plana*)<br>(Baba, 1960) | *Nemopilema nomurai*,<br>*Aurelia coerulea*,<br>*Rhopilema esculentum* | Tang et al. 2021 | 30 - 118 ** |
| ***Coryphella verrucosa***<br>(Sars, 1829) | *Aurelia aurita* | Hernroth & Gröndahl<br>1985 a, b | 200 |
|  | *Cyanea capillata* | Gröndahl & Hernroth<br>1987 |  |
|  | *Aurelia* | Östman 1997 |  |
| (as *Flabellina verrucosa*) | *Aurelia* | Frick 2005 |  |
| ***Cratena pilata***<br>(Gould, 1870)<br>(as *Coryphella* sp.) | *Chrysaora quinquecirrha* | Cargo & Schultz 1967 | 11 polyps in<br>10 min |
|  | *Chrysaora quinquecirrha* | Vogel 1969 |  |
|  | *Chrysaora quinquecirrha* | Oakes & Haven 1971 |  |
| ***Cuthona* sp.**<br>(Alder & Hancock, 1855) | *Chrysaora quinquecirrha* | Soranno 2016 |  |
| ***Dendronotus dalli***<br>(Bergh, 1879) | *Aurelia labiata* | Hoover et al. 2012 |  |
| ***Dendronotus rufus***<br>(O'Donoghue, 1921) | *Aurelia labiata* | Kozloff 1983 |  |
|  | *Aurelia labiata* | Hoover et al. 2012 |  |
| ***Eubranchus* spp.** | *Chrysaora quinquecirrha* | Soranno 2016 |  |
| ***Facelina bostoniensis***<br>(as *Facelina drummondi*)<br>(Couthouy, 1838) | *Aurelia aurita* | Thiel 1962 |  |
| ***Flabellina fusca***<br>(Bergh, 1894) | *Aurelia labiata* | Hoover et al. 2012 |  |
| ***Goniobranchus tinctorius***<br>(as *Chromodoris tinctoria*)<br>(Rüppell & Leuckart, 1830) | *Nemopilema nomurai*,<br>*Aurelia coerulea*,<br>*Rhopilema esculentum* | Tang et al. 2021 |  |
| ***Hermisenda crassicornis***<br>(Eschscholtz, 1831) | *Aurelia aurita* | Keen 1991 | <120 |
|  | *Aurelia labiata* | Hoover et al. 2012 | 8.6 - 31.3<br>polyps h-1 ** |
|  | *Aurelia aurita* | Takao et al. 2014 | 43 - 535 ** |
| ***Pleurobranchaea maculata***<br>(as *Pleurobranchaea novaezealandiae*)<br>(Quoy & Gaimard, 1832) | *Aurelia* sp.1 | Feng et al. 2017 | 393.8 |
|  | *Nemopilema nomurai*,<br>*Aurelia coerulea*,<br>*Rhopilema esculentum* | Tang et al. 2021 | 27 - 78 ** |
| ***Sakuraeolis enosimensis***<br>(Baba, 1930) | *Chrysaora quinquecirrha* | Takao et al. 2014 | 8 - 45 ** |
|  | *Aurelia* sp.1 | Feng et al. 2017 | 26.6 |
| ***Sakuraeolis sakuracea***<br>(Y. Hirano, 1999) | *Chrysaora quinquecirrha* | Takao et al. 2014 | 26 - 131 ** |

### 2.3. Morphological identification of prey nematocysts in the predator’s cerata

*C. militaris* nudibranchs were offered polyps of various Scyphozoa (*R. nomadica*, *C. andromeda*, and *Aurelia* sp.) and Hydrozoa (*A. pluma* and *P. disticha* ) as prey, and the incorporation of nematocysts into the cerata was examined. In order to clear the cerata from previously incorporated nematocysts, and examine the time-line for nematocyst incorporation, nudibranchs were immersed in 5% KCl. As a result, the nudibranchs recoiled and raised their cerata, ejecting a mix of mucus and nematocysts without autotomizing their cerata. However, ten minutes after returning to seawater, the animals resumed normal behavior. While the cerata of freshly collected nudibranchs, at sea, contained hundreds of nematocysts (Fig. 3a), most cerata examined after the treatment were completely empty of nematocysts and in others, between 0 and 3 nematocysts were detected (Fig. 3b).

**Fig. 3:**
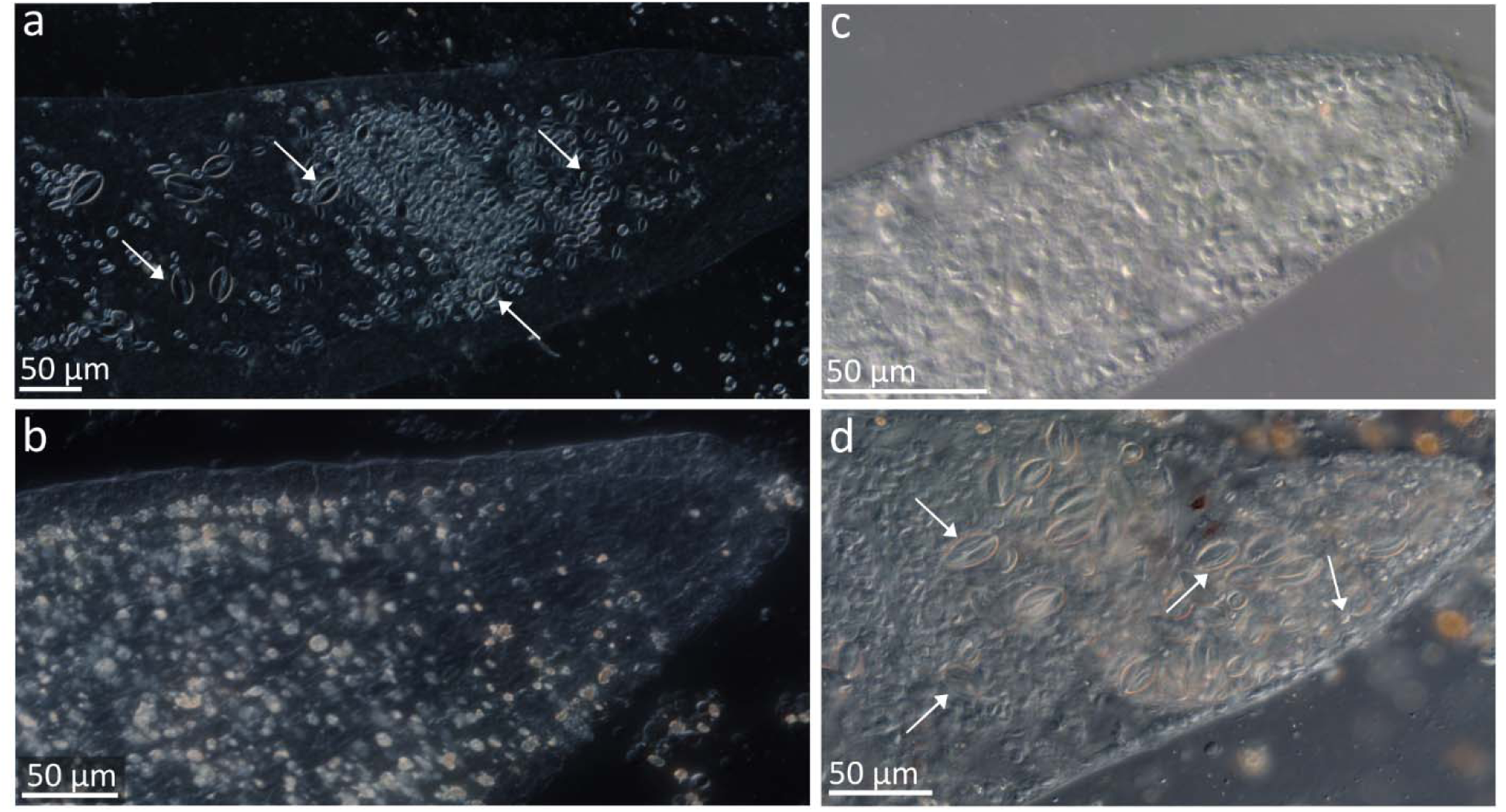
Cerata of *C. militaris* nudibranchs. (a) Naturally feeding (b) After treatment with 5% KCl. (c) Fed with polyps of the scyphozoan, *R. nomadica* on day 12. (d) Fed with the hydrozoan *P. disticha* on day 9.

During the 12 days of the experiment, *C. militaris* nudibranchs preyed on all prey item provided (supp. Table S6). However, the cerata of nudibranchs that fed on the polyps of *R. nomadica*, *C. andromeda*, and *Aurelia* sp., and on the hydrozoan *A. pluma* showed no signs of nematocysts during the experiment at days 3, 6, 9, and 12 (Fig. 3c, presenting a ceras of *C. militaris* feeding on *R. nomadica* ). In contrast, nudibranchs feeding on the hydrozoan, *P. disticha,* incorporated the prey nematocysts into their cerata within 3-6 days, with increasing numbers, temporally, until by day 9 of the experiment, several hundred nematocysts of variou types were concentrated at the tip of each ceras examined (Fig. 3d).

The nematocysts found in the cerata were identified as stenoteles (large, medium, and small), desmonemes, and microbasic mastigophores (large and small) (Fig. 4a), the desmonemes being the most abundant (35.9%), and the large microbasic mastigophores, the least (1.5%) (Table 2). The same types and sizes of nematocysts were identified in the hydrozoan, *P. disticha* , as published by Östman^47^ (Fig. 4b-f) (Table 2). However, the relative abundance of the various nematocyst types was significantly different than that found in the cerata of nudibranchs fed with *P. disticha* (PERMANOVA, R^2^ = 0.56, p < 0.0001). The small stenoteles were the main contributors to the difference between the Hydrozoa and the cerata samples (SIMPER, 39%), but only the desmonemes exhibited similar relative abundances in the cerata and in the Hydrozoa.

**Fig. 4:**
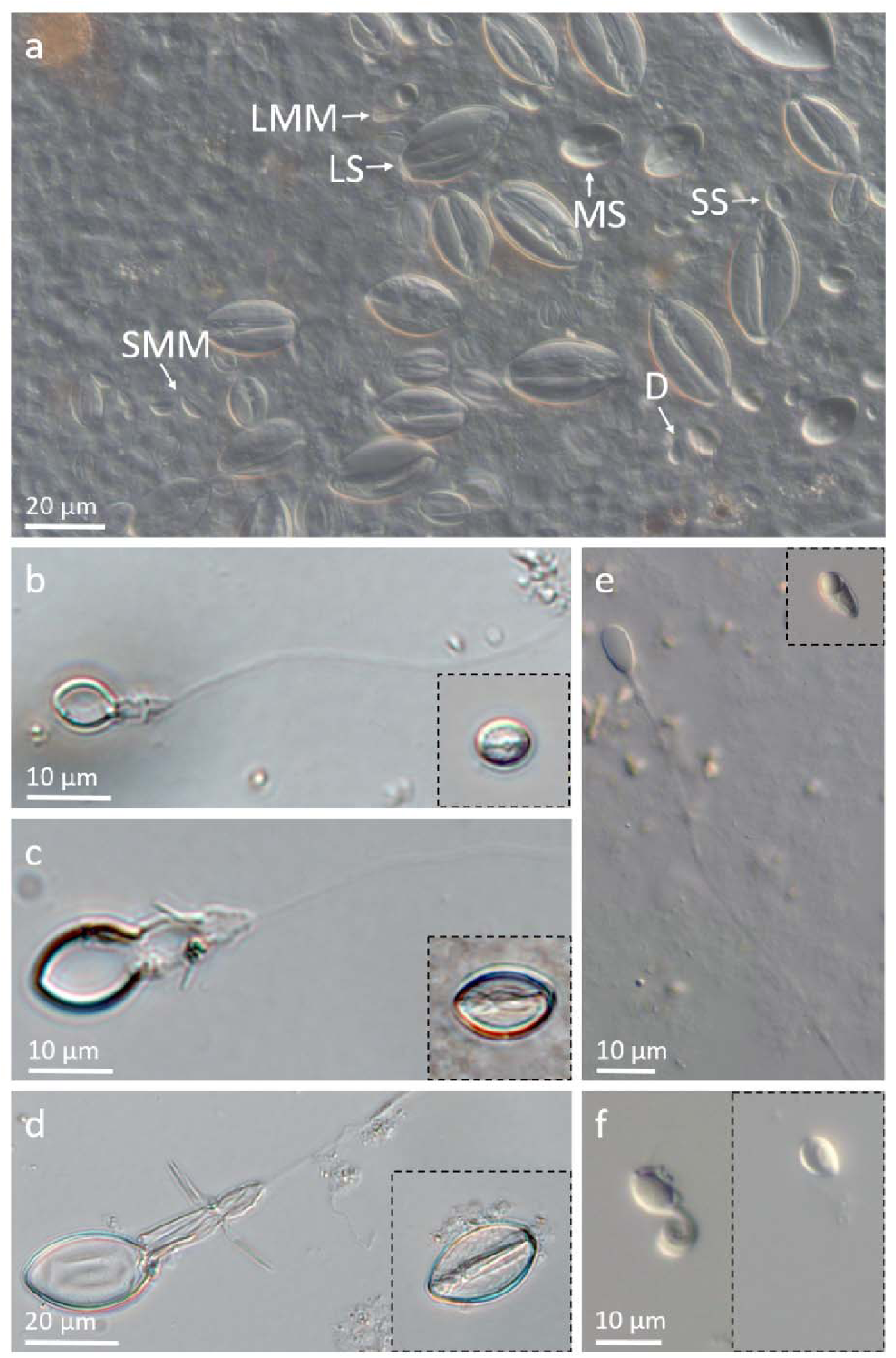
Similar nematocyst types observed in the cerata of *C. militaris* feeding on the Hydrozoa *P. disticha,* and occurring in *P. disticha* (a) Nematocysts in the cerata of *C. militaris* feeding on *P. disticha* (day 9). SS, small stenoteles; MS, medium stenoteles; LS, large stenoteles; LMM, large microbasic mastigophores; SMM, small microbasic mastigophores; D, desmonemes. (b-f) Discharged and intact nematocysts types observed in *P. disticha.* (b-d) Small, medium, and large stenoteles, (e) Microbasic mastigophores, (f) Desmonemes.

**Table 2:**
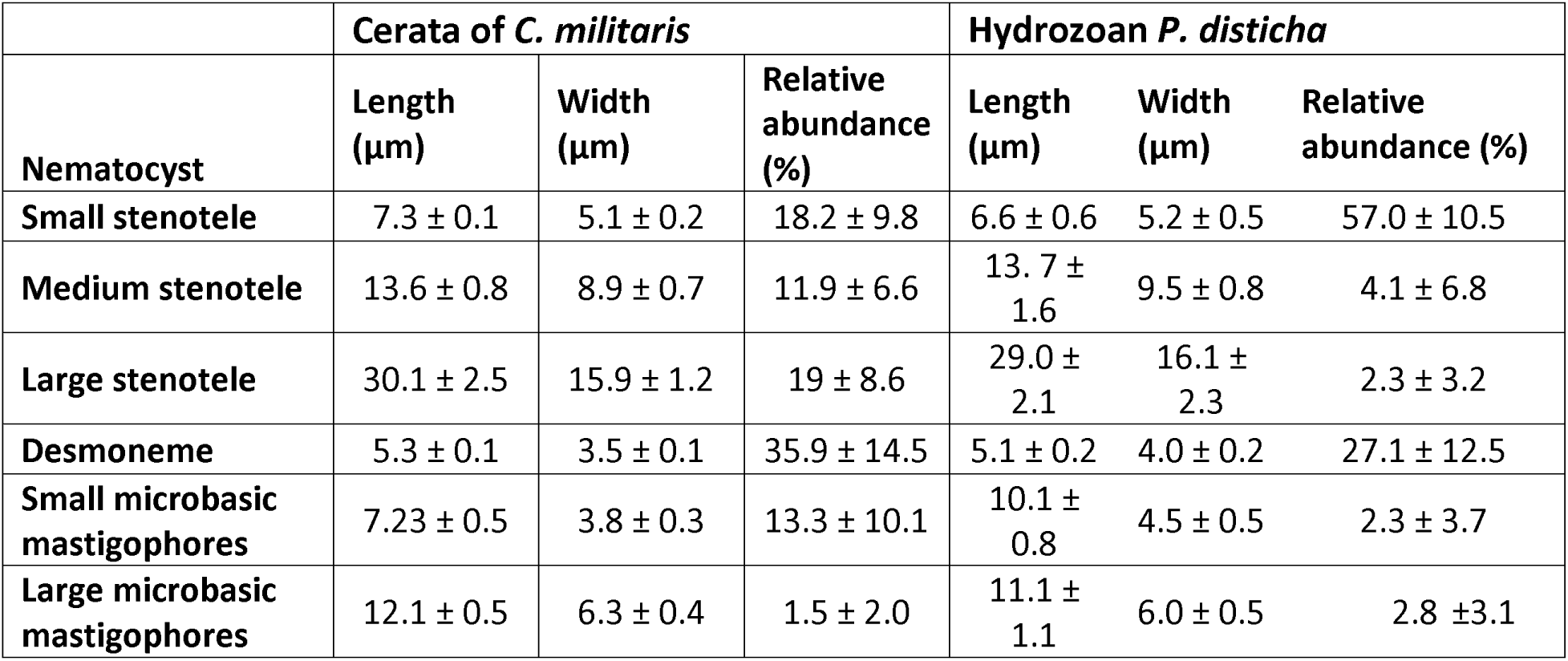
Measurements and relative abundance of the various intact nematocyst types found in the cerata of *C. militaris* nudibranchs after 9 days of preying on the hydrozoan *P. disticha* and in *P. disticha* . Measurements are presented as mean ± SD (n = 10) and relative abundance is presented as percent of >100 nematocysts counted (n = 7).

|  | **Cerata of *C. militaris*** |  |  | **Hydrozoan *P. disticha*** |  |  |
| --- | --- | --- | --- | --- | --- | --- |
| **Nematocyst** | **Length (μm)** | **Width (μm)** | **Relative abundance (%)** | **Length (μm)** | **Width (μm)** | **Relative abundance (%)** |
| **Small stenotele** | 7.3 ± 0.1 | 5.1 ± 0.2 | 18.2 ± 9.8 | 6.6 ± 0.6 | 5.2 ± 0.5 | 57.0 ± 10.5 |
| **Medium stenotele** | 13.6 ± 0.8 | 8.9 ± 0.7 | 11.9 ± 6.6 | 13.7 ± 1.6 | 9.5 ± 0.8 | 4.1 ± 6.8 |
| **Large stenotele** | 30.1 ± 2.5 | 15.9 ± 1.2 | 19 ± 8.6 | 29.0 ± 2.1 | 16.1 ± 2.3 | 2.3 ± 3.2 |
| **Desmoneme** | 5.3 ± 0.1 | 3.5 ± 0.1 | 35.9 ± 14.5 | 5.1 ± 0.2 | 4.0 ± 0.2 | 27.1 ± 12.5 |
| **Small microbasic mastigophores** | 7.23 ± 0.5 | 3.8 ± 0.3 | 13.3 ± 10.1 | 10.1 ± 0.8 | 4.5 ± 0.5 | 2.3 ± 3.7 |
| **Large microbasic mastigophores** | 12.1 ± 0.5 | 6.3 ± 0.4 | 1.5 ± 2.0 | 11.1 ± 1.1 | 6.0 ± 0.5 | 2.8 ± 3.1 |

A comparison between nudibranchs in the feeding experiments and wild-caught animals showed that the cerata of naturally feeding nudibranchs contained various nematocyst types and sizes. Many nematocysts found were small, medium, and large stenoteles (sizes: 5-9x4-8 µm, 13-17x7-14 µm, and 22-26X14-20 µm, respectively), similar in size and shape to the cnidome described above in the cerata of nudibranchs feeding on the hydrozoan *P. disticha.* The most abundant and distinct nematocysts found were bean-shaped microbasic euryteles (ca. 10-16x5-8 µm) with a short shaft, divergent from the lateral capsule axis (Supp. Fig. S7), matching the nematocyst complement of the hydrozoan, *Eudendrium merulum* (Watson, 1985)^48^. Additional nematocysts found in the cerata of naturally feeding *C. militaris* were round stenoteles (9-12x8-11 µm) and small isorhizas (5.5-6.5x4-5 µm). These results show that *C. militaris* selectively incorporates nematocysts into its’ cerata.

## 3. Discussion

The aeolid nudibranch, *C. militaris,* is known to feed on several hydrozoan species^3,25^. Our study examined whether this nudibranch can also prey on scyphozoan polyps and use these as a food source. We show that *C. militaris* preys on various scyphozoan polyps of common eastern Mediterranean species, and that these may serve as an exclusive food source for prolonged periods of time. *C. militaris* exhibited broad predatory behavior, consuming all hydrozoan and scyphozoan polyp species offered, with a maximum intake of 192 *R. nomadica* polyps d^-1^. Feeding on *C. andromeda* polyps facilitated the longest survival period (255 d), whereas nudibranchs feeding on *P. punctata* survived only 40 d. *Aurelia* sp. was the least favorable prey type, showing the lowest predation rate and proportion of polyps consumed (8 ± 10.2 and 27% ± 34%, respectively), while *R. nomadica* was the preferred scyphozoan prey type (44 ± 45.3 and 92.8% ± 11.2% predation rate and proportion of polyps consumed, respectively).

The predation rate of *C. militaris* on jellyfish polyps is comparable with the rates reported for other nudibranch species (see Table 1), suggesting that it can significantly impact polyp populations *in situ* through predation, potentially affecting the size of jellyfish blooms^2,28,33^. However, similarly to other nudibranch species, *C. militaris* did not consume the podocysts of *R. nomadica*^36,41,46,49^. Thus, podocysts may serve as a protective stage, enabling polyp population replenishment even after grazers have reduced the abundances of polyps^50^. *C. militaris* has only recently been introduced into the eastern Mediterranean^23^. Therefore, the predation preferences and long-term ecological effects of this species on jellyfish polyp populations need to be further studied. Given the role of scyphopolyps as pivot contributors to jellyfish blooms^32^, documenting these predator-prey interactions between nudibranchs and scyphopolyps is essential to understand medusan population dynamics.

Ingestive conditioning has been known to affect prey selection in some aeolid nudibranchs, such that they will preferentially prey upon species they have been feeding upon recently^51,52^. *C. militaris* showed no evidence of ingestive conditioning in a similar manner to the behavior observed in *H. crassicornis* ^34^. Although the specific prey of the nudibranchs pre-capture is uncertain, when scyphozoan polyps were offered to freshly-caught nudibranchs, they began feeding on these almost immediately, without an adjustment period. Analysis of cerata from freshly-collected nudibranchs revealed that *C. militaris* exhibits a generalist predatory behavior, such that, in addition to a nematocyst complement consistent with that of the hydrozoans *P. disticha* and *E. merulum,*^47,48^ multiple nematocyst types and sizes were observed.

Nudibranchs that prey on Cnidaria are not immune to nematocyst discharge of the prey, and their response to the sting may vary depending on the prey’s cnidome, size, and the number of nematocysts discharged during the attack. While *C. militaris* was unaffected by hydrozoan nematocysts, the discharge of nematocysts from scyphopolyps led to the retraction of the nudibranch, but did not prevent predation, and nematocyst discharge from the anemone prompted the nudibranchs to retreat. *C. pilata*, a known predator of *Chrysaora quinquecirrha* (Desor, 1848) polyps in Chesapeake Bay, was recorded recoiling upon contact with the polyps’ tentacles^41^. Although stung by the jellyfish polyps, *C. militaris* minimized injury during feeding by approaching the polyps from above, raising the anterior of its body, and engulfing the polyp as a whole (supp. video S2-S4).

Aeolid nudibranchs are known to maintain nematocysts sequestered from their cnidarian prey for defense against predators^1,10,12^. Incorporation of nematocysts is selective and is a function of species, prey choice, and predation pressure^16,18,19,40,53–55^. Our experiments demonstrate that *C. militaris* selectively incorporates nematocysts into its cerata. When provided with scyphozoan polyps and with the hydrozoan *A. pluma*, the nudibranch did not sequester any of the prey nematocysts. In contrast, when provided with the hydrozoan *P. disticha*, nematocysts were detected in the cerata within a few days. This is the first study to report long-term feeding on cnidarian polyps including the ability to feed without incorporation of prey nematocysts. Typically, selectivity in nematocyst incorporation is manifested as a shift in the relative abundance of nematocyst types between the prey and the cerata^14,53,56^. For instance, *C. verrucosa* preying on *Aurelia* sp. polyps, incorporated a much higher ratio of rhopaloids to a-isorhizas, compared to that present in the polyps^40^ and preferentially retained scyphozoan nematocysts over those from other prey species^19^. *C. militaris* displayed a selective avoidance of scyphopolyp nematocysts, and did not retain these nematocysts in its cerata altogether.

Selective incorporation of nematocysts by *C. militaris* was also apparent through the difference in the relative abundance of nematocyst types. As opposed to the nudibranch *C. pilata* (as *Aeolis pilata* ), which incorporated only microbasic mastigophores from the hydrozoan *P. disticha* (as *Pennaria tiarella*)^53^, *C. militaris,* retained all nematocyst types of this hydrozoan in the cerata^47^. However, the relative abundance of the various nematocyst types in the predator was significantly different than that of the prey. This type of selectivity in nematocyst incorporation was reported for many other nudibranchs^14,16,18,19,40,53,56,57^, although in *Berghia stephanieae* (Valdés, 2005) and *Cratena peregrina* (Gmelin, 1791), no direct evidence of selectivity was observed^11,58^.

Variation across nudibranch species in kleptocnidae processes include the types of nematocysts retained, the level of selectivity, nematocyst transfer time, turnover rate, and the quantity of nematocysts stored within each cnidosac^55,59^. For instance, in the nudibranch *B. stephanieae*, new nematocysts appeared in the cerata within 2-4 days of feeding^58^, while in *C. peregrina*, this process took only 2 hours^60^. In our experiments, the first nematocysts appeared in the cerata of *C. militaris* after 3 days. A turnover of nematocyst complement has been documented in *Spurilla neapolitana* (Delle Chiaje, 1841) within six days^14^, while in *H. crassicornis*, it extended over a month^61^. In *C. militaris*, the cerata retained previously acquired nematocysts for over a month when scyphozoan polyps were the exclusive food source.

Intraspecific variation is also substantial: differences in nematocyst composition have been observed in the cerata of individuals from the same species when fed the same diet^54^ and even within different cerata of a single nudibranch^1,60,61^. During our experiments, some cerata contained hundreds of new nematocysts, while others had only just begun to accumulate them. This level of variation raises questions regarding the process and mechanism of nematocyst selection and transport.

Jellyfish blooms have significant ecological, economic, and social impacts worldwide^62^ with scyphopolyps playing a key role in driving these blooms^28,32,63^. However, the location of the polyps of most scyphozoan species *in situ* is unknown. Since this is a major challenge with respect to understanding polyp ecology, many methods (mainly molecular approaches) have been employed to address this problem^21,64–66^. Mills & Miller^67^ were able to identify the prey of the ctenophore *Haeckelia rubra* (Kölliker, 1853) from analysis of nematocysts. Although *C. militaris* does not sequester scyphopolyp nematocysts, the method described here can be used to locate cryptic scyphopolyps *in situ* using other nudibranch species.

Nudibranchs are generally considered specialized feeders, relying on a narrow range of prey species^2^. While *C. militaris* is known to feed on certain hydrozoan species, our findings reveal that it also preys on a variety of scyphozoan polyps, and can use these as a long-term food source. Nudibranchs utilize their cnidarian prey both for nourishment and as a source of nematocysts. Our study shows that *C. militaris* consumes both hydrozoan and scyphozoan polyps, but relies on scyphozoan tissue exclusively for nutrition, selectively incorporating hydrozoan nematocysts for defense. Some nudibranchs have been shown to preferably incorporate scyphozoan nematocysts over the nematocysts of other prey^19^, meaning that these provide adequate protection for nudibranchs against predators. Other nudibranchs rely on cnidarian and non-cnidarian prey for different biological needs: the non-cnidarian supports nourishment and the cnidarian, a source for nematocysts^34^. However, no nudibranch has been reported to utilize Cnidaria only as a food source, discarding its defensive qualities. This type of diversified prey consumption, yet complete selectivity in nematocyst incorporation is unique. Future research exploring the mechanisms underlying nematocyst selection and the ecological consequences of *C. militaris* predation on scyphozoan polyps in the eastern Mediterranean will provide further insights into the role of aeolid nudibranchs in regulating cnidarian populations.

## 4. Methods

### 4.1. Stock cultures of scyphozoan polyps

Stock cultures of *R. nomadica* polyps were produced by laboratory fertilizations of several (4-8) sexually mature male and female jellyfish collected near Mikhmoret, eastern Mediterranean Sea (32°241l231lN 34°521l241lE). To enable planula settlement, glass slides were placed in a fertilization tank with the adult medusae following the protocol described in Dror & Angel ^68^(in press). Polyps of *C. andromeda* have been maintained in the laboratory since their discovery on a rock collected near Jisr az-Zarqa, eastern Mediterranean (32°53’321lN, 34°90’101lE). Polyps of *P. punctata* were produced after collecting planulae from the oral arms of a gravid female medusa collected near Mikhmoret and allowing them to settle onto glass slides. Polyps of an as yet unidentified *Aurelia* sp. were cultivated and maintained in the laboratory from three polyps that appeared in our flowing seawater aquaria. Each species’ polyps were kept in separate containers in a 50 μm filtered flowing seawater aquarium system, maintained at sea surface temperature (SST) ranging annually from 17 to 29 °C. The polyps were fed daily with newly hatched *Artemia salina* nauplii and subjected to a 12-h illumination cycle.

### 4.2. Nudibranch sampling and maintenance

Nudibranchs were collected between 2020-2024 by SCUBA diving near Mikhmoret. The nudibranchs were found on natural rocky (kurkar) reefs at depths of 6-12 m during winter and spring (SST 17-23 °C). Each nudibranch was carefully hand-collected into a 50 ml centrifuge tube and brought to the laboratory for handling. The nudibranchs were kept in 1 L containers of 50 μm filtered flowing seawater, aerated and maintained at SST, subjected to a 12-h illumination cycle.

### 4.3. Short-term predation of nudibranchs on various cnidarian

The short-term feeding experiments were conducted in 200 ml glass bowls of 50 μm filtered seawater with a glass slide (7.5 X 2.6 cm^2^) holding the prey items. Each set of nudibranchs were starved for 24-72 h prior to the experiment to standardize feeding conditions.

Four *C. militaris* (size 1.5-2.5 cm) and four *F. affinis* (size 1-1.5 cm) individuals were each provided with 13-28 *R. nomadica* (Scyphozoa) polyps. In addition, four *C. militaris* individuals (size 1.5-2.5 cm) were each provided with a different cnidarian prey – seven *Exaiptasia diaphana* (Rapp, 1829) (anemone), 14 polyps of *Oculina patagonica* (de Angelis D’Ossat, 1908) (coral), seven polyps of *C. andromeda* (Scyphozoa), six branches of *P. disticha* (Hydrozoa) (Supp. Fig. S8). Another set of three *C. militaris* (size 2-3 cm) were individually introduced into bowls containing three to five 10 cm long tentacles removed from an adult *R. nomadica* medusa (40 cm bell diameter).

The nudibranchs were observed under a dissecting microscope for three hours to study their feeding behavior. The time to the first encounter with a polyp, response to the encounter, and the manner of feeding were recorded. Following this initial close-up examination, the nudibranchs were each placed in an aerated flowing seawater container, maintained at SST, and were allowed to feed for an additional 21 hours (overall 24 h) to record predation rate (number polyps consumed day^-1^), and the predation proportion (number polyps consumed / number polyps provided).

### 4.4. Long-term predation of *C. militaris* on scyphozoan polyps

*C. militaris* nudibranchs were each placed in a 1 L container with flowing 50 µm filtered seawater and with polyps of different scyphozoan species on glass slides. The provided prey polyps included *R. nomadica* , *C. andromeda* , *P. punctata* , and *Aurelia* sp. The polyps were replenished twice weekly, and the predation rate, predation proportion and survival period of *C. militaris* on each species was documented. The experiment lasted for 50 d for the *Aurelia* sp. polyps, and until the nudibranchs died in the other treatments (polyp species). All nudibranchs appeared in good condition during the experiment: feeding continuously, growing, and moving.

### 4.5. Prey nematocysts in nudibranch cerata

#### 4.5.1. Microscopy and photography

Examination of cerata and nematocyst samples was performed by means of Differential Interference Contrast (DIC) microscopy using a Zeiss AXIO Imager.M2 microscope (20X or 40x magnification) and photographed using a digital Axiocam 503 color camera. Measurements of nematocysts were conducted using a Zeiss Zen-Pro 2.5 program.

#### 4.5.2. Induced ejection of nematocysts from *C. militaris* cerata

To remove all the nematocysts from the cerata, nudibranchs were immersed in 5% KCl for 30 s followed, immediately, by three washes in seawater^69^. Two cerata located at the front, and two at the rear of each nudibranch, were carefully removed using a sterile scalpel before and after the treatment. To determine the presence of nematocysts, squash preparations of pre- and post-treatment cerata were examined under a microscope.

#### 4.5.3. Identification of nematocysts in the cerata of *C. militaris*

Cerata (3-5) from each of freshly collected *C. militaris* nudibranchs were removed and stored at -20 °C for identification of nematocysts in naturally feeding nudibranchs. To identify prey nematocysts, the nudibranchs were starved for 96 h, then treated with 5% KCl, and cerata from three nudibranchs were inspected under a microscope to check that all nematocysts were ejected. The experiment was conducted over 12 days, during which the nudibranchs were provided with an excess amount of polyps of the scyphozoan and hydrozoan species as prey, including *R. nomadica* (Scyphozoa, n = 6), *C. andromeda* (Scyphozoa, n = 5), *Aurelia* sp. (Scyphozoa, n = 5), *P. disticha* (Hydrozoa, n = 2), and *Aglaophenia pluma* (Linnaeus, 1758) (Hydrozoa, n = 2). Two-three cerata were removed from each nudibranch on the last day of the experiment (day 12) and stored at -20 °C. In addition, for nudibranchs feeding on *C. andromeda*, *Aurelia* sp. and *P. disticha,* cerata were removed from two nudibranchs on day 3, 6, and 9 of the experiment to determine when the nematocysts appear in the cerata (see supp. Table S9 for sample details). The number of polyps consumed by each nudibranch were counted each time food items were replaced to check that the nudibranchs were actively feeding (supp. Table S6). Squash preparations of the cerata were examined and photographed to check for the incorporation of nematocysts of the various food items in the nudibranch’s cerata. Nematocysts were identified based on Östman^20^ , and the relative abundances of the various nematocyst types in the cerata were documented . For each nematocyst type, width and length of undischarged capsules (n = 10) were measured.

#### 4.5.4. Nematocyst isolation and characterization from the hydrozoan *P. disticha*

Colonies of *P. disticha* were sampled from the sides of the pier in Jaffa port, Israel (32°03?09?N 34°44?57?E). Nematocysts were isolated from 3-5 branches (2-3 cm long) of *P. disticha* (n = 7) by centrifugation of the Hydrozoa in 50% Percoll (Sigma-Aldrich, USA) at 1000 g for 10 min at 4°C. The resulting pellet was re-suspended in DDW and stored at -20 °C. The discharge of nematocysts was achieved by addition of 1 µl of 0.1M NaOH to 3 µl nematocyst suspensions placed on glass slides. For each sample, the relative abundances of the various nematocysts were documented. The description of each nematocyst type included measurements of capsule width and length of discharged and undischarged capsules (n = 10).

### 4.6. Data analysis

Data analysis and graphs were performed using RStudio (version 2023.06.0, R Development Core Team 2023). Analysis of variance (ANOVA) was conducted to check for significant differences in predation rate and proportion of polyps consumed between the various scyphozoan polyps offered as food. All tests were conducted at the α = 0.05 significance level. Assumptions for ANOVA were checked using Levene’s test for homogeneity of variance and a Shapiro-Wilk test for normality. Tukey’s HSD tests were conducted for multiple pairwise post-hoc comparisons of means for the significantly different ANOVA results. Permutational multivariate ANOVA (PERMANOVA) (9999 permutations) were used to test for significant differences in the cnidome of *P. disticha* and the nematocysts found in the cerata of nudibranchs feeding on *P. disticha* . Permutational multivariate analyses of dispersion (PERMDISP) were used to ensure homogeneity of group dispersions and Similarity percentage analyses (SIMPER) were calculated to determine the main contributors to the observed dissimilarity in the cnidomes. Results are presented as % or as mean ± SD.

## Data availability

The datasets generated and analyzed during the current study are available from the authors upon request.

## Supporting information

C. militaris feeding on R. nomadica polyps

C. militaris feeding on C. andromeda polyps

C. militaris feeding on Aurelia polyps

Supplemantary material Dror et al Cmilitaris

## Acknowledgments

We thank R. Yavetz and L. Uziyahu for their valuable technical assistance, the Mevo’ot-Yam students, Applied Marine Biology & Ecology Research (AMBER) laboratory members, Lotan laboratory members, Noga Ratz, Tamar Hamiel, and Amit Comidi for their assistance in nudibranch sampling, Gal Vered for graphic assistance; Dena Restaino for sharing her ideas and knowledge; and Dr. Noa Shenkar and Dr. Miki Kanterovich for the use of their laboratories.

## Author contributions

The research was designed by Hila Dror (HD), Dror Angel (DA), and Tamar Lotan (TL) and executed by HD, DA and various volunteers. Data analyses were conducted by HD. All authors were involved in writing the manuscript and all authors read and approved the final version.

## Additional information Competing interests

The authors declare that they have no conflict of interest.

