## Supplementary material for "Predation on scyphozoan polyps and selective nematocyst incorporation dynamics in the aeolid, *Caloria militaris*": Supplemantary material Dror et al Cmilitaris

### Abstract

Aeolid nudibranchs are known to prey on various cnidarians and incorporate nematocysts from their prey into their cerata (dorsal appendages) for self-defense. Nematocyst incorporation is selective, and is based on species, diet, prey availability, and predation pressure. This study investigated the interactions between two nudibranch species, *Caloria militaris* and *Flabellina affinis*, and scyphozoan polyps from common eastern Mediterranean jellyfish species, including *Rhopilema nomadica*, *Cassiopea andromeda*, *Phyllorhiza punctata*, and *Aurelia* sp. Laboratory experiments were used to assess whether these nudibranchs feed on scyphozoan polyps, and if they incorporate the prey nematocysts into the cerata. Results indicate that while *F. affinis* avoids scyphozoan polyps, *C. militaris* consumes all tested species, and can subsist on them for extended durations (up to 255 days). Predation rates varied among prey species tested, ranging from a mean of 8 to 43.3 polyps day$^{-1}$. However, *C. militaris* does not store scyphozoan nematocysts. These findings contribute to understanding nudibranch feeding ecology and the potential roles these predators play in regulating jellyfish blooms. Identification of scyphozoan nematocysts in the cerata of predator aeolid nudibranchs is discussed as a means to locate cryptic scyphozoan polyps in the marine environment.

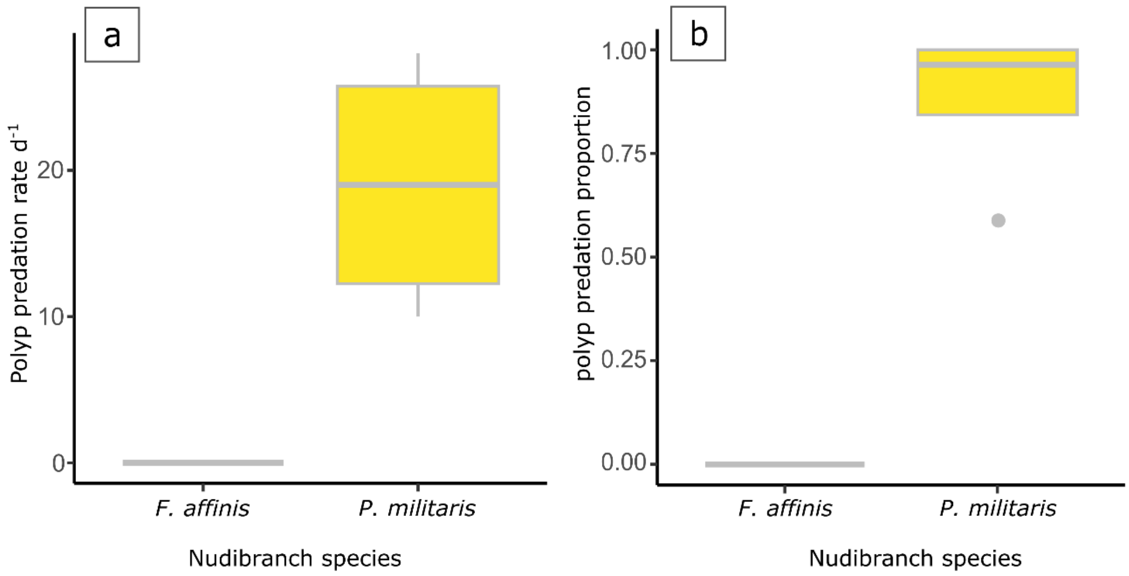


**Supp. Fig. S1**: *R. nomadica* polyps consumed by *F. affinis* (n = 4) and *C. militaris* (n = 4) nudibranchs during 24 h of feeding experiment. (a) predation rate presented as polyps day$^{-1}$ (b) predation proportion (number polyps consumed / number polyps provided).

**Supp. Video S2**: Video capture of *C. militaris* feeding on polyps of *R. nomadica.*

**Supp. Video S3**: Video capture of *C. militaris* feeding on polyps of *C. andromeda.*

**Supp. Video S4**: Video capture of *C. militaris* feeding on polyps of *Aurelia* sp.

**Supp. Table S5**: Summary of the long-term experiment to examine feeding and survival of *C. militaris* nudibranchs on four species of scyphozoan polyps (*R. nomadica, C. andromeda, P. punctata,* and *Aurelia* sp.). * Scyphozoan polyps were replenished twice weekly. ** Survival period for nudibranchs provided with a mix of polyp species is shown as mean ± SD (n = 9). Individual *C. militaris* nudibranchs are identified as "CM" with a number.

| Nudibranch | Size (mm) | Polyp species | # polyps provided feeding $^{-1}$ * | # polyps consumed d$^{-1}$ (mean ± SD) | Survival period (d) ** |
|---|---|---|---|---|---|
| CM 26 | 15 | *R. nomadica* | 15-400 | 44 ± 45.3 | 85 |
| CM 76 | 15 | *R. nomadica* | 4-30 | 2.4 ± 2 | 43 |
| CM 50 | 20 | *C. andromeda* | 29-570 | 34.7 ± 26.6 | 255 |
| CM 51 | 10 | *C. andromeda* | 30-570 | 18.1 ± 13.6 | 146 |
| CM 58 | 10 | *C. andromeda* | 63-280 | 35.7 ± 15.2 | 93 |
| CM 60 | 20 | *C. andromeda* | 55-300 | 47 ± 25.5 | 93 |
| CM 61 | 10 | *P. punctata* | 100-400 | 47.5 ± 19 | 40 |
| CM 62 | 10 | *P. punctata* | 150-400 | 30.1 ± 17.6 | 40 |
| CM 63 | 10 | *P. punctata* | 80-450 | 53.4 ± 36.9 | 40 |
| CM 64 | 30 | *P. punctata* | 230-380 | 40.2 ± 11.6 | 34 |
| CM 107 | 25 | *Aurelia* sp. | 31-157 | 6.5 ± 6.8 | > 50 |
| CM 108 | 15 | *Aurelia* sp. | 26-153 | 13.5 ± 16.6 | > 50 |
| CM 109 | 15 | *Aurelia* sp. | 26-206 | 10.1 ± 9.8 | > 50 |
| CM 110 | 15 | *Aurelia* sp. | 27-180 | 4.4 ± 5.7 | 46 |
| CM 111 | 20 | *Aurelia* sp. | 34-152 | 5.4 ± 6.8 | 46 |

**Supp. Table S6**: Prey species and predation rate (polyps $d^{-1}$) of *C. militaris* nudibranchs during the 12 d feeding experiment (sec 4.5.3). *C. militaris* samples are identified as CM with a number. Presented as mean ± SD (n = 12).

| **Nudibranch** | **Food** | **Predation rate (polyps $d^{-1}$)** |
|---|---|---|
| **CM102** | *C. andromeda* | 10.80 ± 10.84 |
| **CM103** | *C. andromeda* | 6.22 ± 4.31 |
| **CM104** | *C. andromeda* | 5.58 ± 6.71 |
| **CM105** | *C. andromeda* | 1.81 ± 3.79 |
| **CM106** | *C. andromeda* | 9.38 ± 12.34 |
| **CM107** | *Aurelia sp.* | 6.50 ± 6.80 |
| **CM108** | *Aurelia sp.* | 13.54 ± 16.66 |
| **CM109** | *Aurelia sp.* | 10.14 ± 9.84 |
| **CM110** | *Aurelia sp.* | 4.48 ± 5.79 |
| **CM111** | *Aurelia sp.* | 5.41 ± 6.89 |
| **CM102** | *R. nomadica* | 3.93 ± 1.15 |
| **CM103** | *R. nomadica* | 2.37 ± 0.55 |
| **CM107** | *R. nomadica* | 3.27 ± 1.34 |
| **CM108** | *R. nomadica* | 4.27 ± 1.60 |
| **CM113** | *R. nomadica* | 3.39 ± 0.57 |
| **CM114** | *R. nomadica* | 3.83 ± 1.47 |

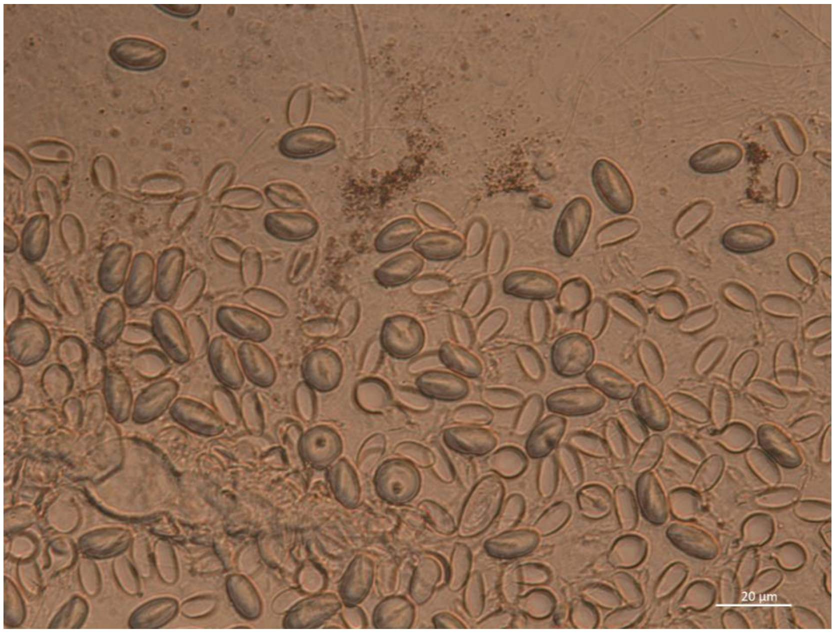


**Supp. Fig S7**: Nematocysts ejected from a squash preparation of a cerasof a naturally feeding *C. militaris* nudibranch showing many discharged and undischarged microbasic euryteles.

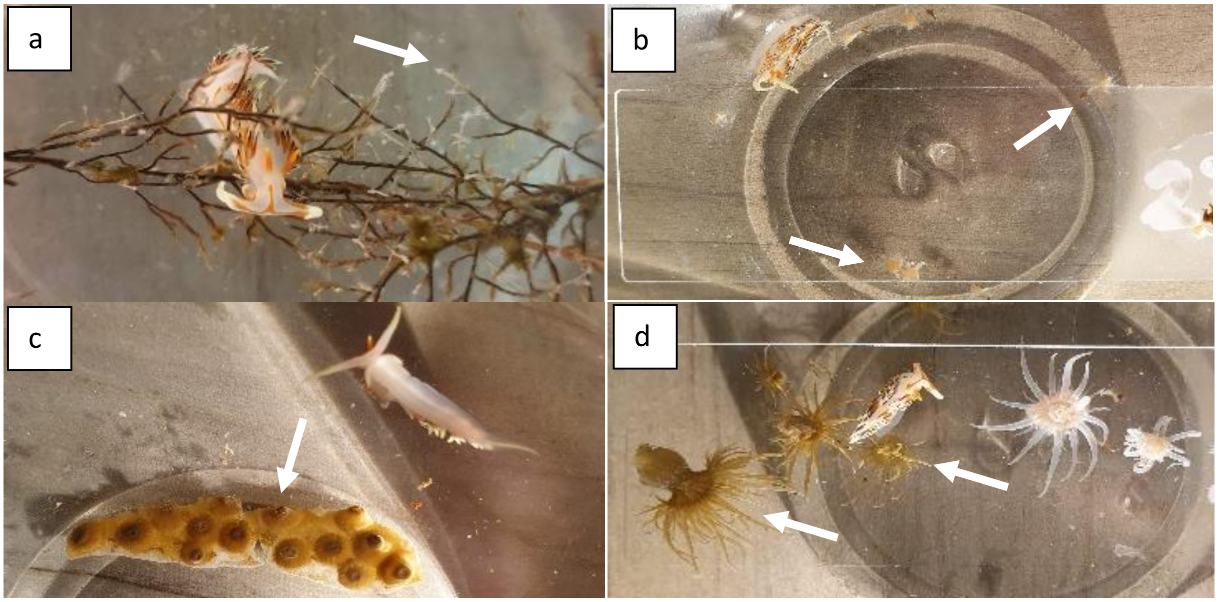


**Supp. Fig. S8**: Experimental glass bowls with various prey species (white arrows) and *C. militaris* nudibranchs. (a) *P. disticha* (b) *C. andromeda* (c) *O. patagonica* (d) *E. diaphana*.

**Supp. Table S9**: Cerata samples taken from *C. militaris* nudibranchs during the 12-day experiment for identification of nematocysts in the cerata. Cerata from naturally feeding (Nat) nudibranchs were sampled, then nudibranchs were treated with 5% KCl, and fed Scyphozoa and Hydrozoa polyps (Cas: *C. andromeda*, Au: *Aurelia* sp., Nom: *R. nomadica*, Ap: *A. pluma*, Pd: *P. Disticha*). Nudibranchs were enumerated CM102 to CM114. Each sample included between 2 and 4 cerata (+ indicates cerata samples).

| | **Day** | | | | |
|---|---|---|---|---|---|
| **Nudibranch-food** | **0 Nat** | **3** | **6** | **9** | **12** |
| **CM102-Cas** | + | + | | + | |
| **CM103-Cas** | + | + | | | + |
| **CM104-Cas** | + | | + | | + |
| **CM105-Cas** | + | | + | | |
| **CM106-Cas** | + | + | | + | |
| **CM107-Au** | + | + | | + | |
| **CM108-Au** | + | + | | | + |
| **CM109-Au** | + | | + | | + |
| **CM110-Au** | + | | + | | |
| **CM111-Au** | + | | | + | |
| **CM112-Ap** | | | | | + |
| **CM113-Ap** | | | | | + |
| **CM102-Nom** | | | | | + |
| **CM103-Nom** | | | | | + |
| **CM107-Nom** | | | | | + |
| **CM108-Nom** | | | | | + |
| **CM113-Nom** | | | | | + |
| **CM114-Nom** | | | | | + |
| **CM113-Pd** | | + | + | + | + |
| **CM114-Pd** | | + | + | + | + |
